# Sensory deprivation triggers phenotypic adjustment of dopaminergic interneurons in the mouse olfactory bulb

**DOI:** 10.1101/2022.07.18.500027

**Authors:** Angelova Alexandra, Tiveron Marie-Catherine, Loizeau Mathieu, Cremer Harold, Platel Jean-Claude

## Abstract

Olfactory sensory activity is a main factor factor controlling intergration and survival of neurons in the olfactory bulb. However, its impact on specific neuronal subtypes is unclear. Using reversible unilateral naris closure in concert with longitudinal in vivo imaging we show here that newborn GABAergic interneurons undergo significant cell death under sensory deprivation. In contrast, dopaminergic OB neurons survive under deprivation, but react with a reversible downregulation of tyrosine hydroxylase expression.

**long abstract:** Neurogenesis persists in the mammalian subventricular zone after birth, producing various populations of olfactory bulb (OB) interneurons. These include GABAergic and mixed dopaminergic/GABAergic double neurotransmitter neurons for the glomerular layer. While olfactory sensory activity is one of the main factors controlling newborn neuron integration, its effect on specific neuronal subtypes is far less clear. Here we use a reversible unilateral deprivation paradigm in combination with longitudinal in vivo imaging to characterize the behavior of newborn glomerular neurons. We find that a substantial fraction of purely GABAergic neurons die after four weeks of sensory deprivation. Tyrosine Hydroxylase (TH) positive dopaminergic/ GABAergic neurons show no signifficant cell death under deprivation, but react with an important decrease in TH expression levels. Importantly, this effect reverses after naris reopening, pointing to a specific adaptation of this neuron population to the level of sensory activity. We conclude that sensory deprivation induces adjustments in the excitation/inhibition balance of the OB implicating cell death and adaptation of neurotransmitter use in specific neuron types.

## Introduction

Neurogenesis persists in the mammalian brain after birth and new neurons are permanently added in the different layers of the mouse olfactory bulb (OB) [1, 2]. Olfactory sensory activity controls the integration of these newborn neurons. Indeed, olfactory sensory deprivation via unilateral naris occlusion (UNO) leads to a large reduction in size of all OB layers and massive cell loss [3-5]. Newborn neurons are particularly affected by UNO, showing increased cell death and decreased dendritic length and spine density [6, 7]. BrdU pulse chase experiments after UNO revealed a decrease in density of newborn periglomerular neurons (PGN) [8].

Newborn PGN represent a heterogeneous population of GABAergic interneurons, distinguishable by the expression of subtype specific markers like Calretinin and Calbindin. Interestingly, a major subpopulation of the GABAergic PGN expresses tyrosine hydroxylase (TH), as well as other enzymes of the catecholamine synthesis pathway, and use dopamine as second neurotransmitter (henceforths called TH-neurons).

While the general impact of UNO on the OB is clear, the consequences on different chemospecific populations is far less well understood. UNO leads to an important loss of TH expression, but only partially of dopamine decarboxylase, another enzyme involved in dopamine synthesis, suggesting an absence of death [9]. More recent studies reported a reduction in numbers of newly generated TH-positive PGNs but not Calretininl□or Calbindinl□expressing PGNs [10] [8] suggesting that only TH-positive PGNs depend on olfactory input for their survival. However, conventional TH-immunostaining did not allow for accurate assessment of their number after naris occlusion [9]. Finally, Sawada and colleagues used a TH-Cre mouse line and found on fixed tissue that 40% of TH PGNs disappear upon UNO [11]. Taken together, there is still lack of consensus about the effect of sensory deprivation on different PGN populations, and specifically in TH positive neurons.

The advent of in vivo two-photon imaging combined with genetic labeling of specific populations allows to unequivocally address this issue. Using this approach, we previously demonstrated that glomerular neuronal populations are differentially affected by UNO since periglomerular glutamatergic interneurons do not undergo cell death under sensory deprivation, while GABAergic neurons do [12]. Here, we followed TH and other GABAergic neurons before, during and after reversible sensory deprivation. We demonstrate that UNO leads to a decrease in TH expression but induces only low level cell death in TH-neurons. In contrast, 15% of GABAergic neurons die during the 4 weeks of sensory deprivation. Reopening the nasal occlusion leads to general arrest of cell death and re-expression of TH-GFP to the initial level in the same neurons.

## Results and Discussion

To visualize and discriminate dopaminergic and non-dopaminergic GABAergic glomerular neurons we performed postnatal electroporation of a Cre expression plasmid into the dorsal ventricular wall of Rosa-RFP /TH-GFP double transgenic mice (Fig. 1A). Four weeks later, after migration in the OB, 20% of RFP positive neurons in the GL were immunopositive for TH. 86% of these RFP+/TH+ neurons were also positive for GFP, showing that TH-GFP represents a reliable proxy of endogenous TH expression in postnatal born OB interneurons (Fig. 1BCD). We also observed that 36% of TH-GFP neurons were negative for TH immunostaining, supporting the notion that TH-GFP expression in TH-neurons precedes the appearance of TH protein [13](Fig. 1C right). In agreement with previous observations [14] we considered the remaining RFP+ PGN as purely GABAergic neurons. Thus, the electroporation approach allowed to discriminate TH-neurons neurons from GABA-only neurons in the OB GL. We then studied the impact of olfactory sensory deprivation on TH protein and TH-GFP levels in the GL. Unilateral naris occlusion (UNO) was achieved by inserting a polyethylene plug into the naris of TH-GFP mice four weeks after CRE-electroporation at P0 [5] (Fig. 1D). Four weeks after UNO, TH-immunofluorescence intensity in the GL was reduced to 18.8% compared to controls (Fig. 1EF). This decrease in TH-protein was paralleled by an 80% reduction in TH-GFP levels (Fig. 1F), confirming previous observations [6] and further validating that TH-GFP reliably reflects protein expression.

**Figure 1:**
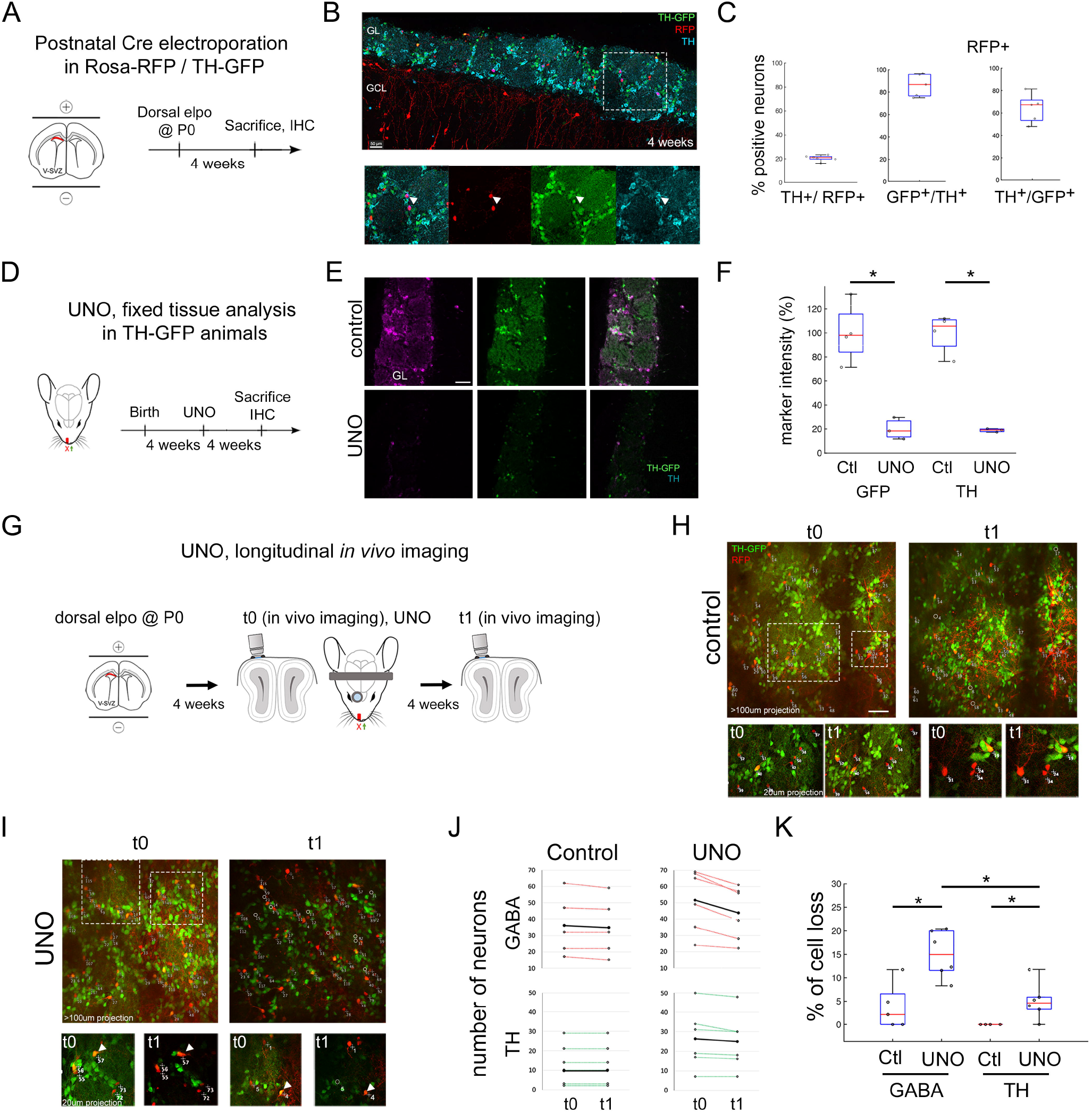
Unilateral naris occlusion reduces immuno-TH and TH-GFP intensity but TH neurons do not disappear. (A) Dorsal electroporation (elpo) of Cre plasmid into postnatal Rosa-RFP/TH-GFP mice is performed at P0 and animals sacrificed at 4 weeks. (B) Four weeks after elpo, RFP+ and RFP+GFP+ interneurons are observed in the glomerular layer of the OB. TH expression of these neurons can be confirmed with anti-TH immunostaining. (C) Around 20% of RFP+ neurons are positive for immuno-TH. 86% of RFP+TH+ neurons are GFP+ and 64% of RFP+GFP+ neurons are TH+. (D) Experimental timeline in UNO experiments. (E,F) TH-GFP and immuno-TH are reduced to 20% of original intensity level after four weeks of UNO. (G) Experimental approach to monitor cell survival *in vivo* after UNO. (H,I) In vivo observation of OB interneurons before (t0) and after UNO (t1) as well as control animals. Circles represent neurons that disappear. Arrows highlight neurons with a decarese of GFP expression. (J) Individual animals and the amount of neurons observed in the course of four weeks in control condition and under UNO. Black line represents the mean number of neurons. Neurons were classified as GABA (RFP+ only) and TH neurons (RFP+GFP+). (K) Quantification of mean number of cell loss in GABA and TH neurons under control and UNO condition. GABA and TH neurons are lost after UNO although significantly less TH neurons than GABA neurons are lost. Statistics: * p<0.05. Scale bars: 50µm (B,E); 40 µm (I).

Next, we used longitudinal *in vivo* 2-photon imaging of postnatally electroporated Rosa-RFP/TH-GFP mice to investigate the impact of sensory deprivation on survival of individually identified TH and GABAergic neurons in the GL. Z-stack recordings of a large field of view (600 × 600 µm), covering the entire depth of the GL, were acquired immediately before (t0, 4 weeks post electroporation) and 4 weeks after UNO (t1, 8 weeks post electroporation; Fig. 1G). Specific glomeruli were easily identifiable based on the circular organization of GFP expressing neurons. Individual RFP+ neurons therein were reliably identified over time, based on their morphology and relative position to neighboring cells (Fig. 1HI). As observed before, we found in control condition a slight expansion of the size of the glomerulus and of the distance between neurons [1] and a shrinkage under UNO [12].

In the absence of UNO, observation of 229 RFP+ GABAergic neurons, including 49 TH-neurons (RFP+GFP+), in 5 mice demonstrated that cell loss in all observed animals was very low, with 3,75 % of the GABAergic neurons (4/180, RFP+GFP-) and no loss of TH-neurons (0/49, RFP+GFP+, Fig. 1HJ). Also, GFP expression in the controls was stable over the observation window (Fig. 1H). Four weeks after UNO, 15% of RFP+ GABAergic neurons were lost (47/310, 6 animals, p<0,05, Fig. 1IJK). TH-neurons showed only a moderate about 5% loss under this condition (n= 9/158 neurons in 6 animals, p<0,05, Fig. 1IJK). However, despite this stability of the TH population after UNO, the striking reduction in TH-GFP expression that was detected immunohistologically was also obvious in vivo (Fig. 1I arrow).

To further explore the plasticity of the glomerular neuronal network we decided to perform reversible UNO in order to investigate how the recovery of sensory activity impacts on cell survival and TH-GFP expression. We performed the same in vivo imaging approach as described above (t0, imaging and UNO), but removed the nasal plug after imaging at t1. The same field of view was then imaged four weeks after reopening (t2, n= 7, Fig. 2A). Control Rosa-RFP/TH-GFP animals (n=6) were submitted to the same labeling and imaging protocol in the absence of UNO. First, we analyzed the presence of GABAergic and TH-neurons over the post-opening period t1-t2. Under control conditions, quantification of RFP cell numbers revealed that overall cell loss at t2 was detected at low levels in all observed animals (GABAergic: 1%, TH: 0.0%, Fig. 2DF). Four weeks after reversal of UNO cell loss was also almost non-detectable in both populations (GABAergic: 1.0%,TH: 0.4%, Fig. 2EF). Thus, the specific impact of UNO on survival of GABAergic neurons was abruptly reversed when sensory information was reinstalled, while the TH population was stable at all observation time points. To assess the evolution of TH expression, we calculated the intensity ratio of GFP over RFP in individual TH-neurons. In control animals, this ratio was unchanged at all three observation time points, as expected (Fig. 2EF; control: n=58 cells, 4 animals UNO: n=57 cells, 4 animals). After UNO, we observed a significant 33% relative loss in GFP intensity in cell bodies (p<0,05, Fig. 2BG). Four weeks after reopening (t2), GFP fluorescence in the identified RFP+GFP+ neurons returned to pre-UNO levels (Fig. 2G and 2C right arrow). Thus, the observed reduction in TH expression in the OB after UNO is not a consequence of loss of this cell population, but represents merely a specific adjustment of this neuron population at the level of neurotransmitter use.

**Figure 2:**
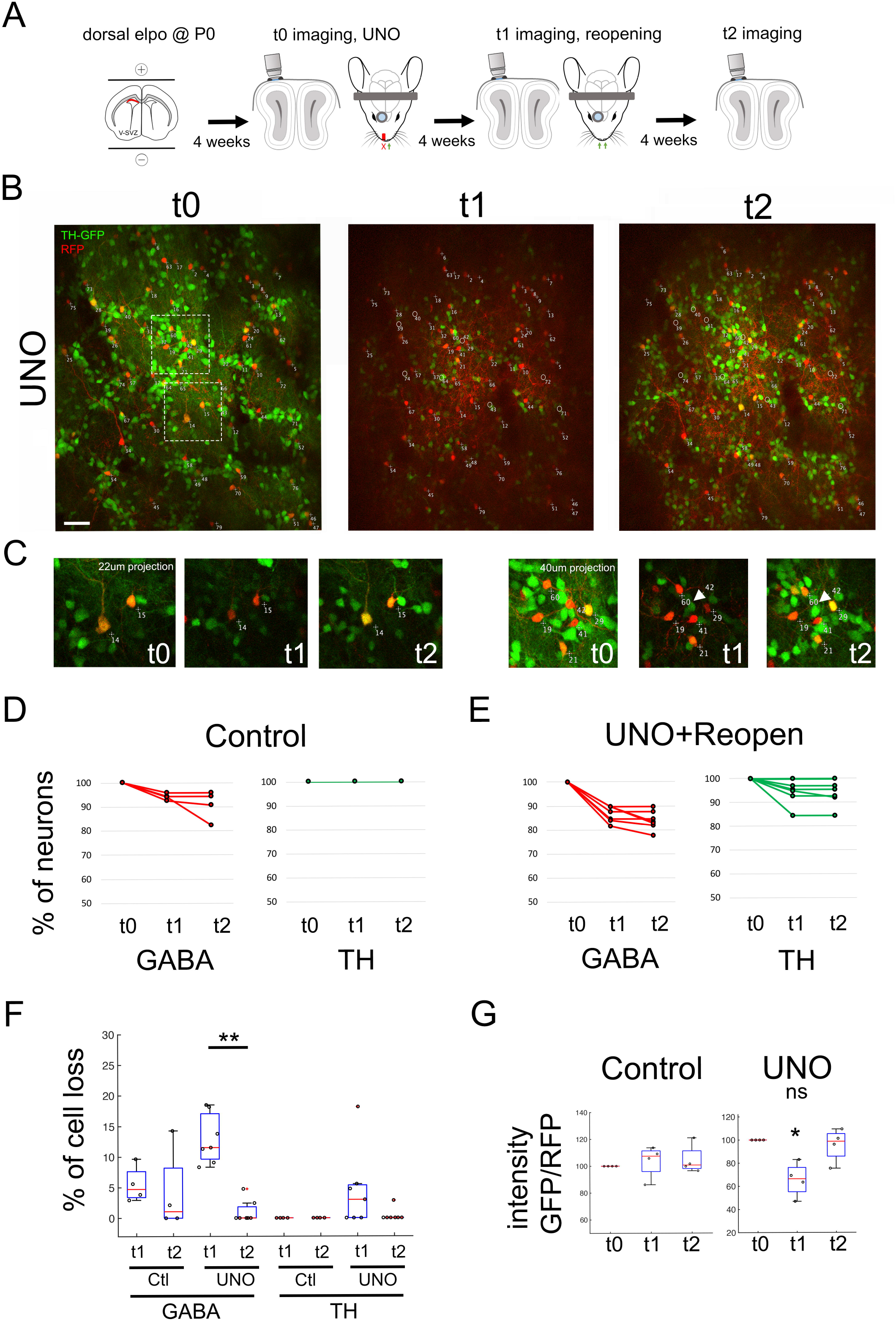
Recovery of sensory activity leads to re-expression of TH in the same neurons. (A) Experimental approach depicting UNO and reopening. (B) *In vivo* observation of the same neurons before UNO (t0), after UNO (t1) and after reopening of naris (t2). Note the strong reduction of GFP in t1 and its re-appearance in t2. (C) close up of surrounded region in B. (DE) Quantification of observed GABA and TH neurons across t0-t1-t2 in control as well as after UNO and reopening within individual animals. (F) Percent of cell loss in GABA and TH neurons across t0-t1-t2 in control and after UNO and reopening. Cell loss in the GABAergic population is significantly reduced after reopening. (G) Ratio of GFP/RFP intensity across t0, t1 and t2. Note the strong reduction of GFP in t1 and its re-appearance in t2. Statistics: * p<0.05; **p<0,01. Scale bar=20µm.

Taken together, we demonstrate that the glomerular neuronal network is capable of a strong reaction to alterations in sensory activity. These adaptations involve plasticity changes in the dopaminergic neurotransmitter phenotype and loss of GABAergic neurons from the network.

## Extended methods

### Animals

All mice were treated according to protocols approved by the French Ethical Committee (#5223–2016042717181477 v2). Mice were group-housed in regular cages under standard conditions, with up to five mice per cage on a 12 hr light–dark cycle. Rosa-RFP mice (Ai14, Rosa26-CAG-tdTomato (Madisen et al., 2010)) and TH-GFP transgenic mice (3846686) were obtained from the Jackson Laboratory and used on a mixed C57Bl6/CD1 background. All experiments were performed on males and females.

### Postnatal electroporation

*In vivo* electroporation was performed as previously described (Boutin et al., 2008). Briefly, P0-P1 pups were anesthetized by hypothermia and 2 μl of pCAG-CRE plasmid (#13775 Addgene, at 4μg/μl) were injected in the lateral ventricle. Animals were subjected to five 95V electrical pulses (50 ms, separated by 950 ms intervals) using the CUY21 edit device (Nepagene, Chiba, Japan) and 10 mm tweezer electrodes (CUY650P10, Nepagene) coated with conductive gel (Control Graphique Medical, France). Electrical pulses were applied to target the dorsal V-SVZ. Electroporated animals were then reanimated in a 37°C incubator before returning to the mother.

### Cranial window implantation

Implantation of a cranial window was performed as previously described (Drew et al., 2010) with minor modifications. Briefly, 4 week old mice were anaesthetized by intraperitoneal (ip.) injection of ketamine/ xylazine (125/12.5 mg/kg). Dexaméthasone (0.2 mg/kg) and buprenorphine (0.3 mg/mL) were injected subcutaneously and lidocaine was applied locally onto the skull. The pinch withdrawal reflex was monitored throughout the surgery, and additional anesthesia was applied if needed. Carprofen (5 mg/kg) was injected ip. after the surgery. A steel head-fixation bar was added attached to the skull with dental cement (Superbond, GACD). A 2.5 × 1.5 mm piece of skull overlying the OB was carefully removed using a sterile scalpel blade. Great care was taken not to damage the dura. A thin layer of low toxicity silicon adhesive (Kwik-Sil) was applied over the craniotomy and covered with a round 3 mm coverslip. The craniotomy was sealed with superglue and dental cement (Superbond, GACD). The first time point (t0) of our microscopic observation was performed after surgery on these anesthetized mice.

### Unilateral naris occlusion

For olfactory sensory deprivation, a polyethylene tube was inserted (BD Intramedic, PE50, 3mm long) into one naris and sealed with cyanoacrylate glue (Cummings et al., 1997). Efficiency of occlusion was checked the following day and before each imaging session by the absence of air bubbles after application of a water drop on the occluded naris. Occlusion lasted for 4 weeks. At the end of the experiment immunostaining against tyrosine hydroxylase was performed to further confirm the efficiency of the occlusion. For reversible occlusions, mice were anesthetized after 4 weeks of occlusion and the polyethylene tube was carefully removed. If the quality of the cranial window has declined due to regrowth of connective tissue and bone, a new surgery was performed to reopen the window and allow for subsequent imaging.

### *In vivo* two-photon imaging

We used a Zeiss LSM 7MP two-photon microscope modified to allow animal positioning under a 20X water immersion objective (1.0 NA, 1.7mm wd) and coupled to a femtosecond pulsed infrared tunable laser (Mai-Tai, SpectraPhysics). After two-photon excitation, epifluorescence signals were collected and separated by dichroic mirrors and filters on 4 independent non-descanned detectors (NDD). Images were acquired using an excitation wavelength of 950 nm. GFP was collected at 500-550. RFP was first collected between 605-678. In addition, we collected an additional RFP signal between 560-590 that was voluntarily saturated to allow a better identification of subcellular structures like dendrites. In general, image acquisition lasted about 10 min. Mice could potentially move on a treadmill during imaging, but rarely did so. The imaging window was centered on the dorsal surface of the OB. The whole PG layer was imaged (around 150 µm depth).

For longitudinal observation, we used the same principle as applied in Platel et al. 2019. In short, the same field of view was localized based on the geometric motifs of groups of neurons and specific morphological features of individual cells. Images of 606×606 µm were acquired at 0.59 µm/pixel resolution in the xy dimension and 2 µm/frame in the z dimension to a maximal depth of 200 µm.

### Chronic *in vivo* imaging analysis

Quantitative analyses were performed on raw image stacks using FIJI software (Schindelin et al., 2012). All neurons identified on the first image were assigned a number using ImageJ overlay. Based on morphology and relative position each neuron was individually numbered and tracked on the successive imaging sessions. Results were summarized in Microsoft Excel. Occasionally neurons located at the border of an image were placed outside of the imaged field in one of the following sessions. These cells were excluded from further analyses. Animals showing an evident degradation of the imaging window were excluded from further imaging sessions.

### GFP fluorescence evolution

To determine the effect of UNO on GFP intensity and to be independent of laser power, we used one wavelength to excite both GFP and RFP and performed ratio calculation as follow: we drew region of interest over the cell bodies of GFP positive neurons. We extracted both the fluorescence intensity of the red channel and of the green channel at t0, t1 and t2. We then performed a ratio green/red for each time point for each cell. We then normalized the ratio for each cell to t0 ratio. Finally we averaged this values per animal (4 control animals and 4 UNO animals).

### Immunohistochemistry and image analysis

For histological analysis, mice were deeply anaesthetized with an xylazine/ketamine overdose. Intracardiac perfusion was performed with 4% paraformaldehyde in PBS. The brain was collected and incubated overnight in the same fixative at 4°C. Sections were cut at 50 µm using a microtome (Microm). Standard immunostaining protocols were performed on coronal free floating sections. Briefly, sections were rinsed in PBS and incubated in a blocking buffer (10% fetal bovine serum (FBS), 0.3% Triton X-100 in PBS) for one hour. Subsequently, sections were incubated in primary antibody solution (5% FBS, 0.1% Triton X-100 in PBS (PBST)) and Tyrosine Hydroxylase antibody (chicken Ig, AVES; 1:1000) overnight at 4°C. The following day, sections were rinsed 3 times in PBS and incubated with species-appropriate secondary antibody in PBST for 2h at RT using gentle rocking. Alexa Fluor-conjugated secondary antibodies were purchased from Life Technologies. Nuclear counterstain HOECHST 33258 (Invitrogen, 1:2000) was added before sections were washed in PBS and mounted using Mowiol as a mounting medium. Optical images were taken either using a fluorescence microscope (Axioplan2, ApoTome system, Zeiss) or a laser confocal scanning microscope (LSM510 or LSM780, Zeiss, Germany). Image analysis was performed using FIJI software (Schindelin et al., 2012). All experiments and quantifications were performed blindly to experimental groups. For figure 1E and F, ROIs were drawn over the whole glomerular layer and the mean intensity of GFP or TH compared between control and UNO animals (4 control and 3 UNO animals)

### Statistical analyses

All data are presented as mean ± s.e.m. In box plot representation, center line represents the median; box limits, upper and lower quartiles; whiskers, outliers). Statistical comparisons were performed using R. Two-sample Fisher-Pitmann permutation test was used as a two-tailed test.

